# Approximate Bayesian Computation applied to time series of population genetic data disentangles rapid genetic changes and demographic variations in a pathogen population

**DOI:** 10.1101/2022.10.24.513475

**Authors:** Méline Saubin, Aurélien Tellier, Solenn Stoeckel, Axelle Andrieux, Fabien Halkett

**Affiliations:** Université de Lorraine, INRAE, IAM, F-54000 Nancy, France; Professorship for Population Genetics, Technical University of Munich, Freising, Germany; INRAE, Agrocampus Ouest, Université de Rennes, IGEPP, F-35653 Le Rheu, France

## Abstract

Adaptation can occur at remarkably short timescales in natural populations, leading to drastic changes in phenotypes and genotype frequencies over a few generations only. The inference of demographic parameters can allow understanding how evolutionary forces interact and shape the genetic trajectories of populations during rapid adaptation. Here we propose a new Approximate Bayesian Computation (ABC) framework that couples a forward and individual-based model with temporal genetic data to disentangle genetic changes and demographic variations in a case of rapid adaptation. We test the accuracy of our inferential framework and evaluate the benefit of considering the full trajectory compared to few time samples. Theoretical investigations demonstrate high accuracy in both model and parameter estimations, even if a strong thinning is applied to time series data. Then, we apply our ABC inferential framework to empirical data describing the population genetics changes of the poplar rust pathogen following a major event of resistance overcoming. We successfully estimate key demographic and genetic parameters, including the proportion of resistant hosts deployed in the landscape and the level of standing genetic variation from which selection occurred. Inferred values are in accordance with our empirical knowledge of this biological system. This new inferential framework, which contrasts with coalescent-based ABC analyses, is promising for a better understanding of evolutionary trajectories of populations subjected to rapid adaptation.

## 1 Introduction

Adaptation can occur at remarkably short timescales in natural populations, leading to drastic changes in genotype frequencies and phenotypes over a few generations only (Buffalo and Coop, 2019). This rapid pace of adaptation has motivated the use of temporal data to understand neutral (Prout, 1954; Wallace, 1956; Nei and Tajima, 1981; Pollak, 1983; Mueller et al., 1985b; Waples, 1989; Wang and Whitlock, 2003) and under-selection genetic evolution over time (Dobzhansky, 1943; Fisher and Ford, 1947; Kettlewell, 1958, 1961; Mueller et al., 1985a). However, such time series studies remain rare compared to the amount of work focusing on one contemporary sample to trace back its genetic history (Buffalo and Coop, 2019, 2020; Pavinato et al., 2021). Temporal data allow to track the changes in allele frequency through time, and therefore lead to a better understanding of evolutionary processes (Dehasque et al., 2020; Feder et al., 2021; Saubin et al., 2022a). In cases of rapid adaptation especially, the resulting genetic changes may be transient and require time sampling around the selection event to be highlighted (Saubin et al., 2022b).

The inference of demographic parameters can allow understanding how evolutionary forces (especially genetic drift and selection) interact and shape the genetic trajectories of populations during rapid adaptation (Bergland et al., 2014; Živković et al., 2019). However, the difficulties in obtaining the likelihood of models including both demography, selection, and genetic drift (Pavinato et al., 2021) lead to consider alternative approaches relying on simulations (Bazin et al., 2010; Laval et al., 2019). With the advent of computational approaches, Approximate Bayesian Computation (ABC) has become a standard approach for genetic analyses aimed at tracing back the evolutionary history of populations (Rosenberg and Nordborg, 2002; Bazin et al., 2010; Cornuet et al., 2010; Estoup and Guillemaud, 2010). These methods, coupled with a coalescent simulator, allow inferring evolutionary scenarios and demographic parameters, by modelling the genealogy of the samples (Whitlock and Barton, 1997). The coalescent is computationally very efficient but relies on strong modelling assumptions (Rosenberg and Nordborg, 2002). As such, coalescent-based ABC approaches suffer from two limitations. First, there is a gap between the models developed so far and the actual complexity of biological scenarios. Different coalescent models have been developed to account for fluctuating population sizes (Sjödin et al., 2005) or partially clonal species (Orive, 1993; Hart-field, 2021). Yet these specificities has been considered in isolation. To date, few models integrate both rapid demographic fluctuations and complex life cycles (Tellier and Lemaire, 2014). Second, coalescent-based inference methods are powerful only when coalescent events occur at the same time scale as the process being considered, such as past demographic events. This means that demographic events happening too fast compared to the coalescent time scale (in units of the effective population size *N_e_*) cannot be inferred. For example, under antagonistic coevolution, changes in the demography of host and pathogen are too fast for a coalescent analysis, but tracking allele frequencies and nucleotide diversity forward-in-time is informative about this process (Živković et al., 2019). With a few exceptions (*e.g*. De Mita and Siol, 2012; Foll et al., 2015), time samples are not considered in such frameworks. Rapid adaptation processes are associated with transient genetic changes that can be well illuminated by considering step-by-step modelling algorithms forward-in-time. The study of the rapid adaptation of pathogens would therefore benefit from being considered with forward models (Foll et al., 2015). Yet they consider so far relatively simple demography. In the case of more complex scenarios, the first step is to decipher how a selective event shape population structure at neutral loci (Hoban et al., 2016). Here we propose a new ABC framework that couples a forward and individual-based model with temporal genetic data to disentangle genetic changes and demographic variations in a case of rapid adaptation.

Understanding and inferring the evolutionary trajectories of populations is of major interest to evolutionary biologists but it can also have practical applications for population management. This is especially the case in agriculture where the need to control pathogen populations is paramount (Parat et al., 2016). Pathogens can induce disease outbreaks devastating human-managed ecosystems (Anderson et al., 2004; Tobin, 2015; Savary et al., 2019). Understanding the evolution of pathogens is therefore crucial for developing effective disease management strategies (Bonneaud and Longdon, 2020). The rapid adaptation of pathogen populations (McDonald and Linde, 2002; Saubin et al., 2021) and the high stochasticity in pathogen evolutionary trajectories (Parsons et al., 2018) make this endeavour extremely difficult.

The rapid adaptation of pathogen populations in agrosystems results from the tremendous selection pressures exerted by modern agricultural practices (Zhan et al., 2015; Stukenbrock and McDonald, 2008). To counter plant disease outbreaks, breeders develop resistant plant genotypes. These genetic resistances are most often deployed across large spatial scales (Zhan et al., 2015; Rimbaud et al., 2021). These agricultural practices break eco-evolutionary feedbacks that maintain the polymorphism observed in natural host-parasite systems (Stukenbrock and McDonald, 2008; Brown and Tellier, 2011). As such, these practices weaken the sustainability of plant genetic resistances in favouring the emergence and spread of virulent (*i.e*. resistance-adapted) pathogens (Rimbaud et al., 2021; Saubin et al., 2022a). Therefore, the outcome of plant genetic resistance deployment is often a resistance overcoming event, *i.e*. the failure of the host plant to remain resistant to the pathogen. This results in the spread of virulent pathogens on resistant hosts (Johnson, 1984; Pink and Puddephat, 1999; Brown and Tellier, 2011; Burdon et al., 2016). On the pathogen side, an event of resistance overcoming can translate into a strong selective sweep with the intense and unidirectional selection causing drastic demographic changes for the pathogen population (Burdon et al., 2016; Persoons et al., 2017; Saubin et al., 2021). Such rapid adaptation can lead to specific temporal genetic signatures at neutral loci, depending on the evolutionary scenario ruling the change in population sizes (Saubin et al., 2022b).

Here, we propose to use ABC based on temporal genetic evolution to unravel the evolutionary scenarios following rapid and contemporaneous adaptation. We apply our inferential framework to time series data and evaluate the added value of considering the full trajectory following forward simulations compared to few time samples. We test the accuracy of several temporal sampling designs in inferring model parameter values. Last, we apply our ABC inferential framework to empirical data describing the population genetics changes of the poplar rust pathogen following a major event of resistance overcoming (Persoons et al., 2017; Louet et al., 2021).

## 2 Materials and methods

### 2.1 Simulation model

The rapid adaptation we model is a resistance overcoming event that is monitored through time. We use an individual-based, forward-time and non-spatial demogenetic model, designed for diploid individuals (Saubin et al., 2021, 2022b). This model couples population dynamics and population genetics to follow the evolutionary trajectory of different genotypes at the selected locus and at neutral genetic markers. The model is implemented in Python (version 3.7, van Rossum, 1995) and Numpy (Harris et al., 2020). We consider life cycles commonly found in temperate pathogen species, with seasonal variation in reproductive mode. These pathogens switch from clonal reproduction during the epidemic phase to sexual reproduction occurring once a year, in winter (Agrios, 2005). This general life cycle is adjusted in two variations to take into account that the sexual reproduction takes place either on the same host plant as for the epidemic phase or on an alternate host (usually a different species). These life cycles are named with or without host alternation, respectively. Without host alternation, the model represents the evolution in time of a population of pathogens on two static host compartments: a resistant compartment (R) and a susceptible compartment (S). Each compartment has a fixed carrying capacity for the pathogen population, *K_R_* and *K_S_*, respectively. With host alternation, the alternate host compartment (A) is added, to account for the sexual reproduction occurring on another species. This static compartment is assumed to be larger than the two other compartments, with a fixed carrying capacity *K_A_*. In the following, we refer to the three compartments as S (susceptible hosts), R (resistant hosts) and A (alternate hosts). The successive steps modelled through generations are presented Figure 1, and detailed in Saubin et al. (2021). We simulate the evolution at neutral loci and at a selected locus responsible for the virulence (qualitative trait) of pathogen individuals. Evolution at neutral loci is set to suit classical population genetic analyses based on microsatellite markers: 23 loci with a mutation rate of 10^*−*3^ and a maximum of 20 allelic states.

**Figure 1:**
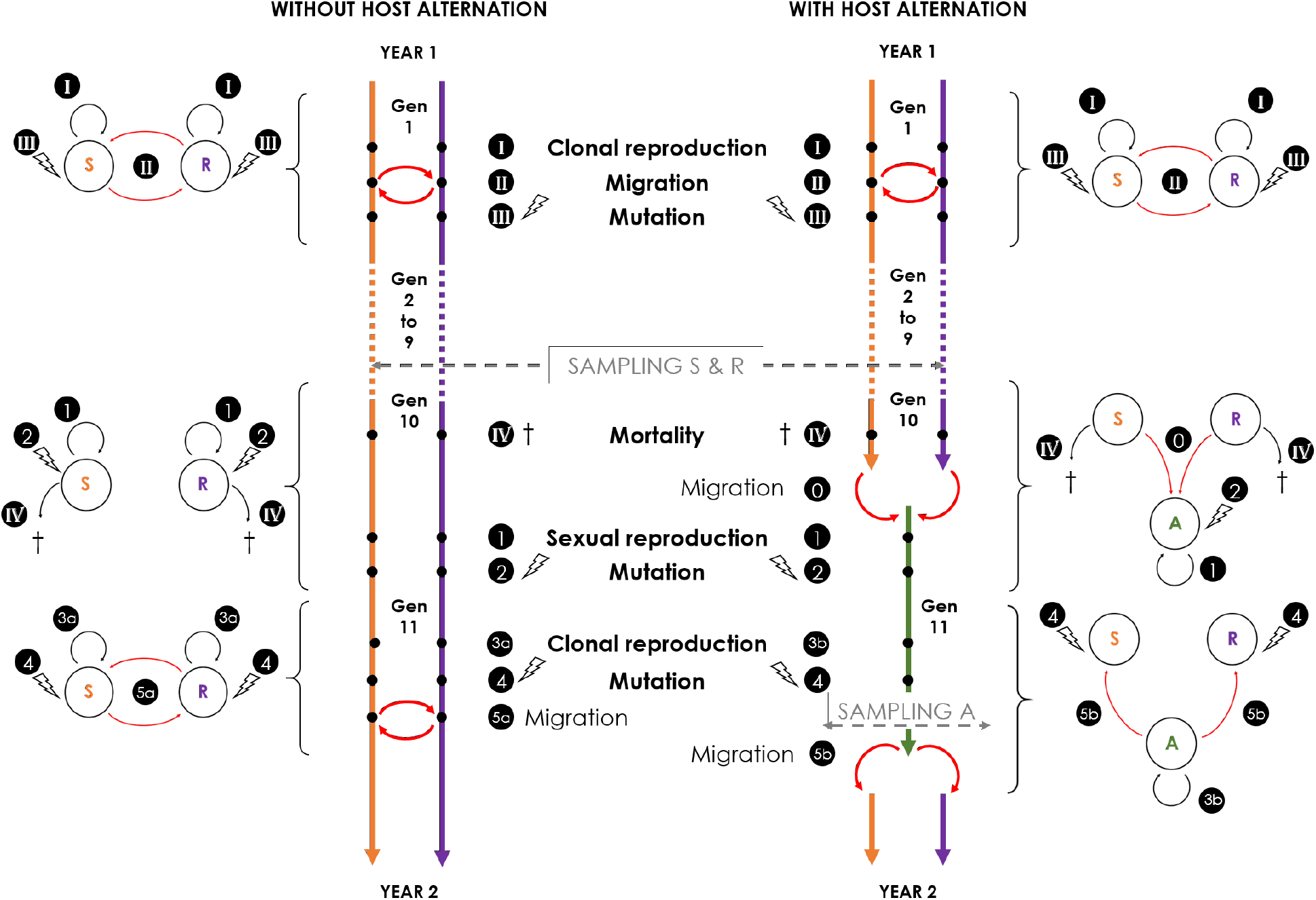
Modelling steps for each simulated year with the three S, R, and A host compartments. Each year is composed of 11 generations. During the clonal phase (generation 1 to 9), each generation is composed of three steps identical between both life cycles: (I) clonal reproduction; (II) migration of a proportion *mig* of each population between R and S; (III) mutation at all neutral markers with a mutation rate of 10^*−*3^. At the end of the clonal phase, the pathogen overwinter as a dormant stage and is subjected to (IV) mortality of a proportion *τ* of each population. Then, the sexual phase (generations 10 and 11) differs depending on the life cycle: (0) represents the migration of all individuals from R and S towards A; (1) sexual reproduction; (2) and (4) mutation of all neutral markers with a mutation rate of 10^*−*3^; (3a) and (3b) clonal reproduction; (5a) migration of a proportion *mig* of each population between R and S; (5b) migration of all individuals from A towards R and S. A sampling takes place every year at the end of generation 9 on S and R and at generation 11 before the migration event (5b) on A.

Each simulation starts with genotypes randomly drawn from the 20 possible alleles followed by a burn-in period with a population of constant size *N* = *K_S_*. During the burn-in period, the pathogen population only evolves on S. To reach the genetic drift and mutation equilibrium from the initial state, the burn-in period is set to 2*N* generations. We build a random simulation design of 150,000 simulations, with the following parameter values drawn randomly from defined prior distributions (Table 1): 1) the proportion *f_avr_* of virulent alleles introduced after the burn-in period in the pathogen population, reflecting initial standing genetic variation; 2) the life cycle of the pathogen (*Cycle*, with or without host alternation); 3) the migration rate (*mig*) of the pathogen population; 4) the growth rate (*r*) of the pathogen population; 5) the annual mortality rate of the pathogen population before the sexual reproduction (*τ*); 6) the cumulative carrying capacity of S and R (*K* = *K_R_* + *K_S_*); and 7) the proportional size of R 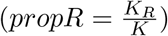.

**Table 1:** Input parameters and their range of variations for the random simulation design run for 400 generations.

| Variable | Description | Distribution | Interval |
| --- | --- | --- | --- |
| $f_{avr}$ | Initial frequency of the virulent allele in the pathogen population | Log-uniform | $[\log(0.0005); \log(0.3)]$ |
| $Cycle$ | Life cycle of the pathogen | Binomial | Without or with host alternation, probability 0.5 |
| $mig$ | Migration rate between R and S | Log-uniform | $[\log(0.001); \log(0.2)]$ |
| $r$ | Growth rate of the pathogen population | Uniform | $[1.1; 2]$ |
| $\tau$ | Mortality rate during the annual migration | Uniform | $[0.5; 1[$ |
| $K$ | Cumulative carrying capacity of susceptible and resistant hosts | Uniform | $[1,000; 20,000]$ |
| $propR$ | Proportion of resistant hosts in the landscape | Uniform | $]0.01; 0.99[$ |

Each simulation is run for 400 generations (with 11 generations per year). We retain only simulations for which the frequency of virulent allele in the population on S at the end of the simulation is above the threshold of 60%. This threshold is chosen to ensure that only simulations effectively leading to the resistant host invasion and resistance overcoming are kept. Of the 150,000 simulations produced from the random simulation design, 106,058 lead to resistance overcoming and are used for the following analyses (50,547 with host alternation, and 55,511 without host alternation).

### 2.2 Summary statistics

Sampling represents a random draw of *n* individuals from a host compartment, with the sample size *n* = 30. The sampling occurs each year at the end of generation 9 on S and R, and additionally - for the life cycle with host alternation - at generation 11 on A before the redistribution to S and R (Figure 1). Sampled individuals are not removed (sampling with replacement) so that their genotypes could contribute to subsequent generations.

To summarise the observed and simulated datasets, we compute the following statistics for each sample of individuals: (i) the proportion of virulent individuals (*P_V ir_*); (ii) the genotypic diversity estimated by Pareto’s *β* (*β_P_*); (iii) the proportion of unique genotypes (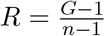 with *G* the number of unique genotypes and *n* the number of sampled individuals); (iv) the genetic diversity estimated by Simpson’s index (*S*); (v) the genetic diversity estimated by Shannon-Wiener’s index (*S*); (vi) the mean expected heterozygosity over all loci (*MH_E_*); (vii) the variance of the expected heterozygosity over all loci (*V H_E_*); (viii) the mean number of alleles per locus (*MLA*); (ix) the multilocus index of linkage disequilibrium 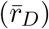; the population differentiation index between populations on R and S sampled at the same generation (*F_ST_ R−S*); (x) the population differentiation index between the initial population (on S after burn-in) and the sampled population (*TF_ST_*) (Table 2). All statistics are implemented along with the model. Summary statistics are calculated for each population, independently of the sampled compartment (S, R or A), except for *P_V ir_* which is not recorded for populations sampled on R (because all individuals living on the resistant host are virulent), and for *F_ST_ R − S* which is calculated from populations on S and R from the same generation. In the following analyses, summary statistics from different compartments are considered as distinct summary statistics. Because of the temporal sampling, each summary statistic is recorded for multiple generations, which constitutes a time series of summary statistics.

**Table 2:**
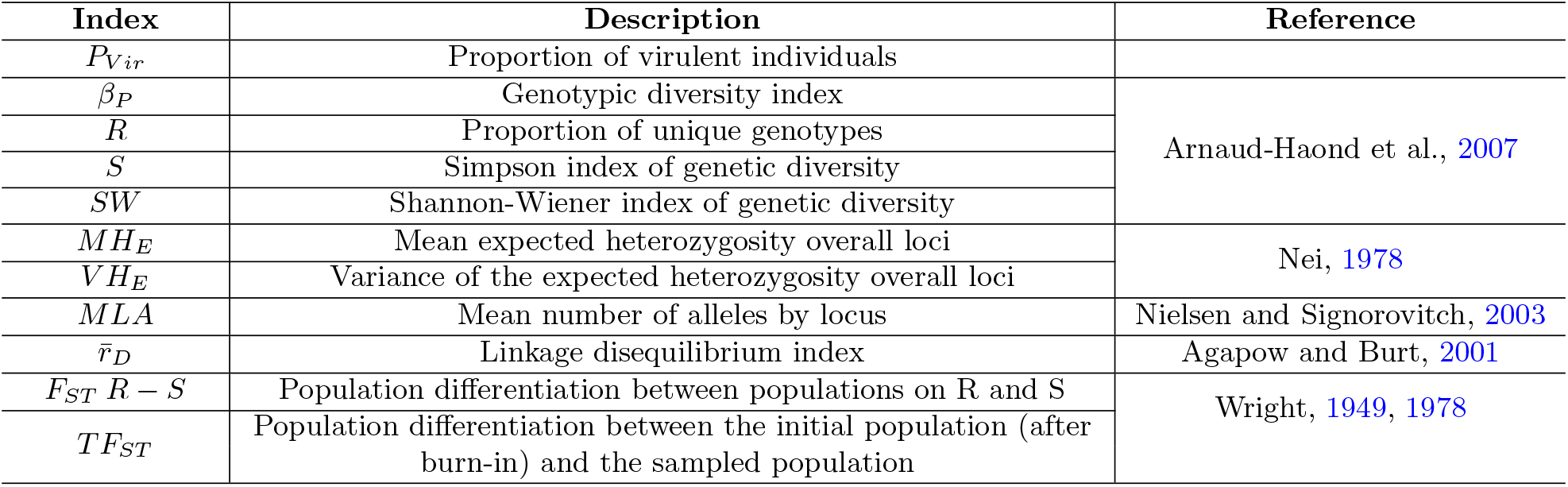
Description of population genetics indices used as summary statistics in the ABC analyses.

Two methods are used to perform the analyses: 1) Summary statistics from different generations are considered as distinct summary statistics and hence all generations along the time series are taken into account (hereafter referred to as ‘Complete summary statistics’); 2) We extract the mean, variance, minimum and maximum values along each time series, and considered these four values as the distinct set of summary statistics for each population genetic index (hereafter referred to as ‘Wrap-up summary statistics’).

### 2.3 Approximate Bayesian Computation

We use the R package Abc (Csilléry et al., 2012) to perform an ABC analysis from the simulated data. This analysis involves two steps: model choice and parameter estimation. The model choice aims to distinguish between the two simulated life cycles, with and without host alternation. The parameter estimation aims to estimate each of the six quantitative parameters of the chosen simulation model (Table 1).

The model choice procedure is based on a weighted multinomial logistic regression (Fagundes et al., 2007) computed on 1% of the simulated datasets for which *ϵ*, the Euclidean distance between the summary statistics of the observed dataset and the simulated datasets, is the smallest (Beaumont et al., 2002). Bayes factors are calculated as the ratio of the posterior probabilities for the tested models (Kass and Raftery, 1995).

For the parameter estimation procedure, we use the 50,547 simulations obtained under the model with host alternation. We estimate posterior distributions (mode and 95% confidence intervals) of each parameter by applying the neural networks regression method (Blum and François, 2010) implemented in the R package Abc based on the 1% of simulations closest to the observed data.

#### Cross-validation on simulated data

To assess the validity of the method, cross-validations are performed at both steps of the analysis. A leave-one-out cross-validation for the model choice is performed on 500 randomly drawn simulations from each of the two models: with and without host alternation. From the Bayes factors obtained, the probabilities to re-estimate the true model are calculated for each life cycle (*i.e*. the model that is indeed used for the randomly drawn simulation). Additionally, we perform Principal Component Analyses (PCA) from the values of summary statistics obtained from the two life cycles. We display envelopes containing 95% of the simulations. A leave-one-out cross-validation for the parameter estimation is performed on 200 randomly drawn simulations from the 50,547 simulations under the life cycle with host alternation.

#### Sampling scheme

In the theoretical case presented in the results, we calculate summary statistics based on the maximum information available. We name it the full simulation design, as we consider all time samples, that is populations sampled once a year for 36 years on all host compartments (S and R for the model choice; S, R and A for the parameter estimation). To assess the impact of the sampling scheme on inference accuracy, we consider two types of rarefaction affecting the time series and the range of compartments considered (Appendix A).

### 2.4 Case study: overcoming of poplar rust resistance RMlp7

Here we apply our ABC inference framework to a documented case of resistance overcoming by a diploid plant pathogen responsible for the poplar rust disease, *Melampsora larici-populina* (Basidiomycota, Pucciniales). This pathogenic fungus is a host-alternating species. Its life cycle consists of an annual sexual reproduction on larch needles in early Spring, followed by clonal multiplications on poplar leaves from spring to autumn (Duplessis et al., 2021; Louet et al., 2021). Its sexual reproduction is obligatory in temperate climates because of the annual fall of poplar leaves (Xhaard, 2011). To control poplar rust disease, several resistant poplars have been selected and planted widely in Western Europe over years. However, the intensive and monocultural plantations combined with the host species being perennial (Gérard et al., 2006) makes poplars particularly vulnerable to the adaptation of the pathogen. This leads to regular events of resistance overcoming, so that all known resistance types have now been overcome. (Pinon and Frey, 2005; Louet et al., 2021). The most damaging resistance overcoming by *M. larici-populina* occurred in 1994 with the adaptation of the pathogen to the resistance RMlp7 carried at that time by a vast majority of cultivated poplar trees. The adaptation of the pathogen to the resistance RMlp7 resulted from the association of two alterations (a nonsynonymous mutation and a complete deletion) at the candidate locus *AvrMlp*7, from standing genetic variation Louet et al., 2021). This led to a rapid invasion, in less than four years, of adapted pathogens across Western Europe, including France (Barr`es et al., 2008; Xhaard et al., 2011; Persoons et al., 2017), causing drastic epidemics (Pinon et al., 1998; Pinon and Frey, 2005). This rapid adaptation event strongly shaped the resulting genetic structure of *M. larici-populina* populations (Xhaard et al., 2011; Persoons et al., 2017).

Poplar rust individuals were sampled before, during, and after this resistance overcoming event and correspond to the samples analysed in the population genetics study of Louet et al., 2021. Each sampled individual was genotyped with 20 microsatellite markers, and population genetics indices are calculated from the model developed above. Two versions of this temporal sampling are used. The first data set is composed of poplar rust samples from the same geographic location (Amance, France), collected from susceptible poplars (S) at the end of the epidemic season and from larch (A) after the sexual reproduction. Twenty-eight populations were sampled between 1989 and 2021 (22 on S, and 6 on A). Sampling size ranges from 5 to 58 individuals (samples with less than 5 individuals were removed from the analyses). The second data set is composed of poplar rust samples from a broader geographic region (Grand Est region, France), collected from susceptible and resistant poplars (S and R) at the end of the epidemic season, and from larch (A) after the sexual reproduction. This second data set includes all individuals from the first data set. Thirty-six populations were sampled between 1988 and 2021 (26 on S, 4 on S and 6 on A). The sampling size ranges from 5 to 79 individuals (samples with less than 5 individuals were removed from the analyses).

For this case study, cross-validation procedures are performed as described above, based on simulated data with the same sampling schemes as the two biological data sets. For the two data sets, the accuracy of the model choice and parameter estimation are compared depending on the choice of summary statistics (Complete summary statistics, or Wrap-up summary statistics). Model choice and parameter estimations are then performed as described above on the empirical data sets, and the posterior distribution is obtained for each model parameter.

For the model choice, we check the goodness-of-fit by computing the *P − value* to test the fit of the observed data to each model. For each model, we reject the null hypothesis model if *P − value* < 0.01.

## 3 Results

### 3.1 Model choice and parameter estimations for the full simulation design

We first evaluate the accuracy of our ABC inference under the ideal case with the maximum data available: populations are sampled every generation from all host compartments.

The cross-validation procedure for the model choice highlights a strong accuracy of model choice when Wrap-up summary statistics are used (Table 3). Conversely, the model identifiability is weaker for Complete summary statistics. However, whatever the summary statistics used, there is a strong overlap in the outcome of simulations realised under the two life cycles (Figure 2). Hence only small areas of parameter values allow to truly discriminate between the two models.

**Table 3:** Accuracy of model selection for the full simulation design depending on the type of summary statistics considered. The model choice procedure is based on leave-one-out cross-validations with a weighted multinomial logistic regression computed with tolerance parameter set at 0.01, for 500 replicates.

| Summary statistics | Probability to find the true model |  |
| --- | --- | --- |
|  | Without host alternation | With host alternation |
| Complete | 0.76 | 0.80 |
| Wrap-up | 1.00 | 0.99 |

**Figure 2:**
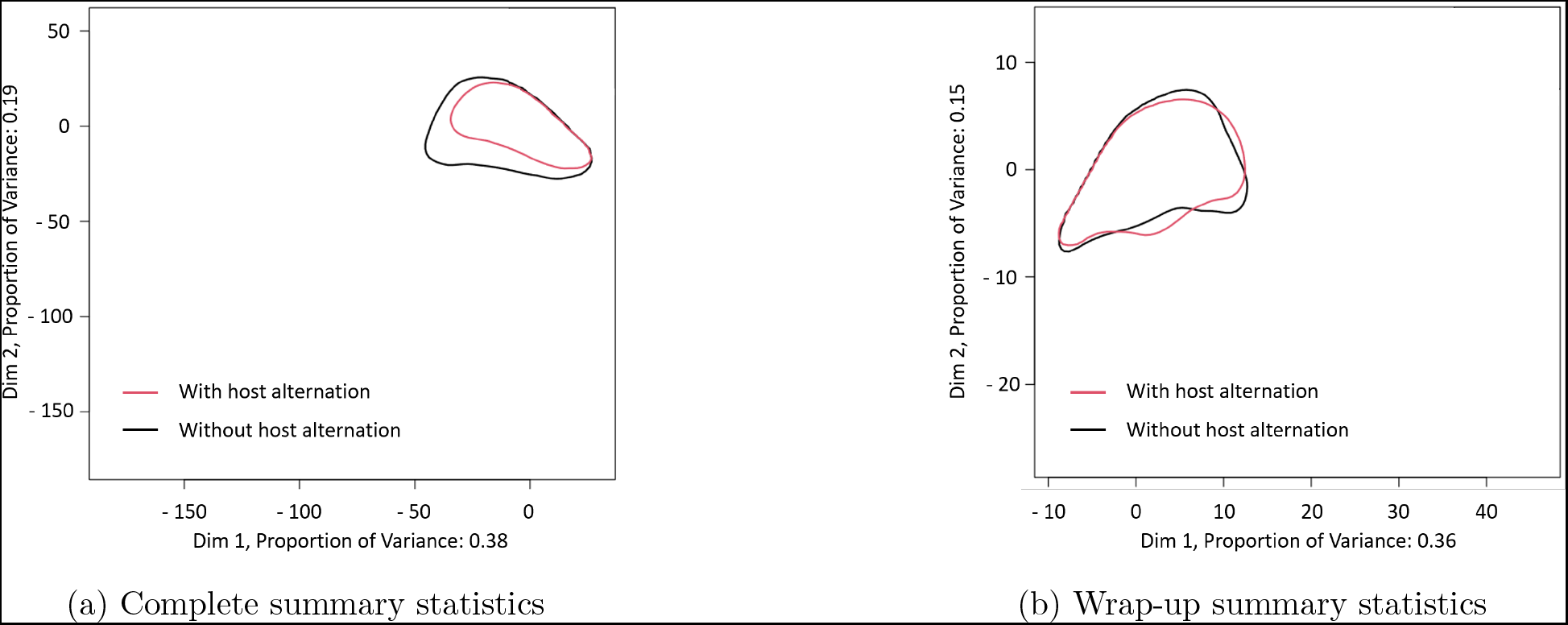
Principal component analyses (95% envelope) of simulations for each model: with and without host alternation, depending on the summary statistics considered. PCA analyses were based on the full simulation design.

The cross-validation procedure for parameter estimation highlights a strong discrepancy among parameters. Some parameters are very well estimated, including *propR*, *K*, *f_avr_* and *r*, ranked in a decreasing order of identifiability (Table 4, Figure 3). The worst parameter to estimate is *mig*, irrespective of the simulated value (Figure 3). Last, *τ* is badly estimated, but with increasing confidence as its true value increases (Figure 3).

**Table 4:** Accuracy of parameter estimation for the full simulation design depending on the summary statistics considered. Data represent r-squared of the linear regressions between simulated and estimated parameters. The parameters identifiability procedure is based on a leave-one-out cross-validation with the neural networks regression method and tolerance parameter set at 0.01, for 200 replicates.

| Summary statistics | $propR$ | $mig$ | $f_{avr}$ | $r$ | $K$ | $\tau$ |
| --- | --- | --- | --- | --- | --- | --- |
| Complete | 0.91 | 0.41 | 0.68 | 0.73 | 0.81 | 0.58 |
| Wrap-up | 0.94 | 0.60 | 0.86 | 0.85 | 0.92 | 0.79 |

**Figure 3:**
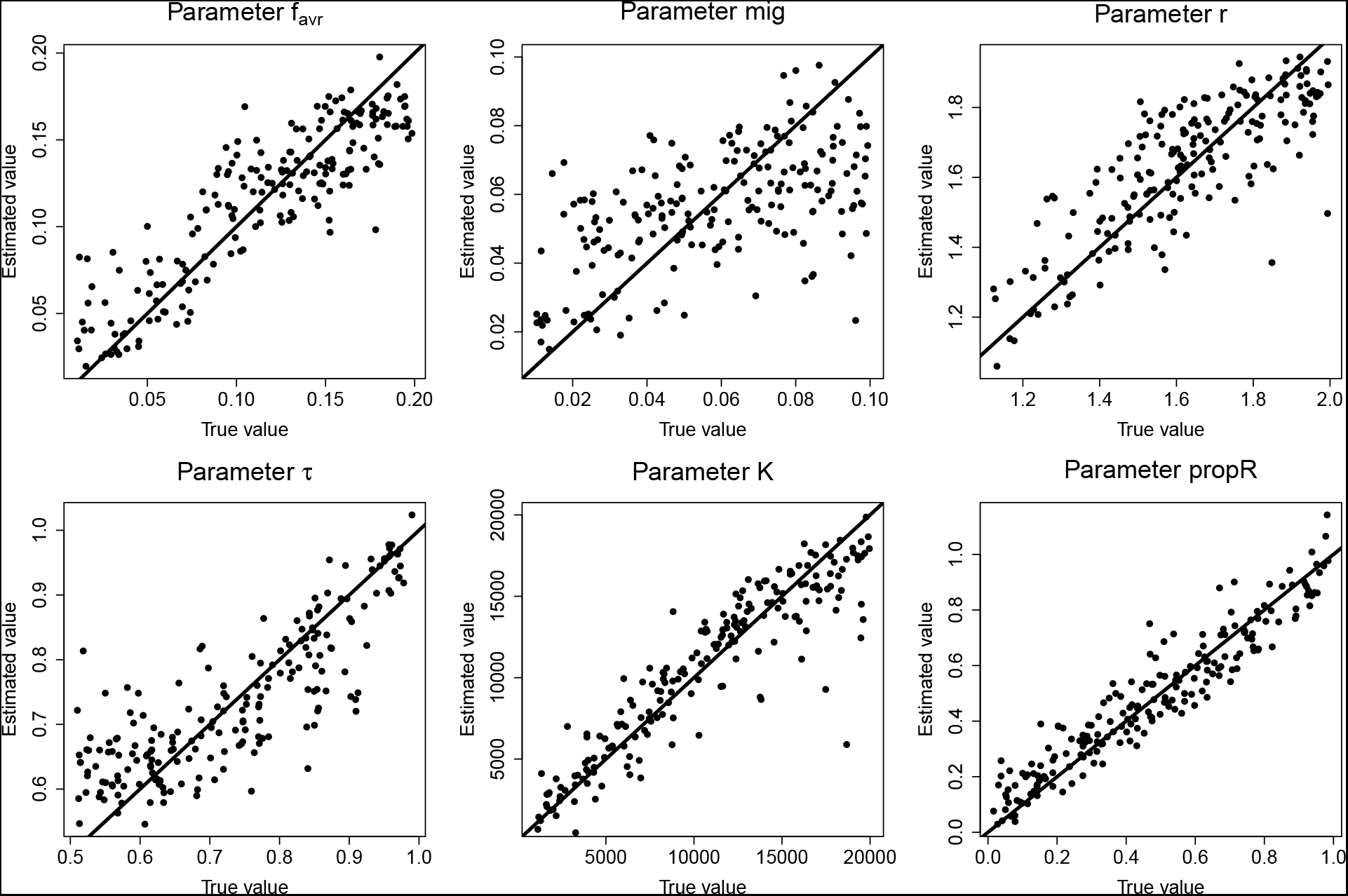
Practical identifiability of parameter estimation for Wrap-up summary statistics. Each point represents the parameter estimation (‘Estimated’ value) depending on the real parameter (‘True’ value). Each graph regroups the results of 200 replicates. Straight lines correspond to the first bisector.

### 3.2 Model choice and parameter estimations when rarefaction is applied

For both cross-validations, sampling every five years gives similar results than considering the full time series of samplings. Reducing the range of sampled compartments also provides fairly good estimates for the model choice and parameter estimations. This result holds even if the sampling focuses on S only. Overall, the ABC accuracy is higher for Wrap-up summary statistics than for Complete summary statistics. Last, the accuracy of the model choice drops drastically if only the first and last time points are considered (Table S1) and the parameter estimations are less accurate, especially for *r* and *τ*. However *propR*, *f_avr_* and *K* are still well estimated, regardless of the rarefaction (Table S2).

### 3.3 Case study: Approximate Bayesian Computations applied to the poplar rust resistance overcoming

In this section, we apply our ABC framework to the two empirical data sets describing the rapid evolution of the genetic structure of a pathogen population following resistance overcoming.

#### 3.3.1 Cross-validations applied to the sampling schemes of empirical data sets

We first perform the cross-validation procedures with sampling schemes corresponding to the two data sets (Amance location and Grand Est region). For both data sets, the cross-validation of model choice shows weak identifiability of life cycles (Table 5). The identifiability of the life cycle with host alternation is slightly better than without host alternation but still limited. The model is more identifiable with the sampling scheme of the data set from the Grand Est region, which represents more time points and both S and R compartments, contrary to the data set from Amance location sampled on S only. As for the full simulation design, the accuracy of parameter estimations is particularly high for parameters *propR* and *K*, regardless of the data set (Table 6). The estimation of parameters *f_avr_* is more mitigated but still quite good (*r − squared* > 0.7 for the two data sets), especially with Wrap-up summary statistics. Conversely, *mig*, *r* and *τ* are less accurately estimated, especially with Complete summary statistics. Overall, as for the full simulation design, the ABC performs better using Wrap-up summary statistics than Complete summary statistics.

**Table 5:**
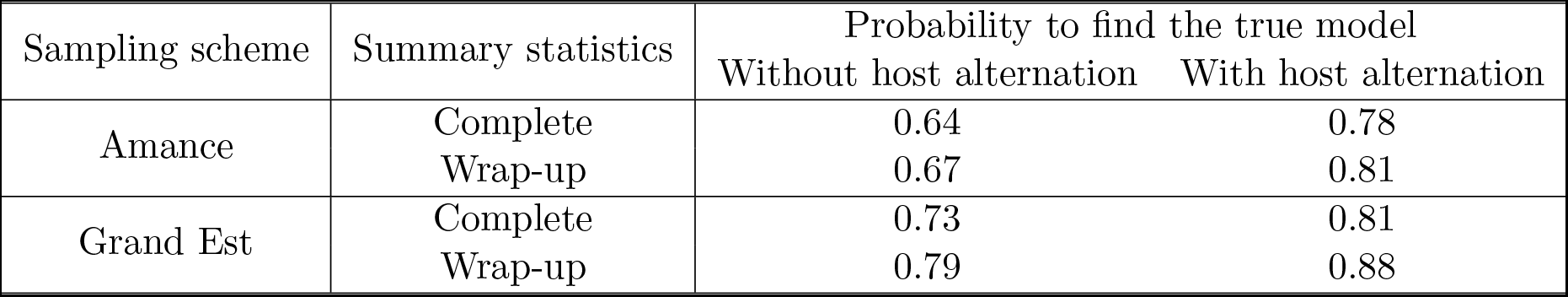
Accuracy of model selection on simulated populations on S and R, depending on the summary statistics considered. The sampling schemes of simulation data sets match those of the two the empirical data sets: Amance location and Grand Est region. The model choice procedure is based on leave-one-out cross-validations with a weighted multinomial logistic regression computed with tolerance parameter set at 0.01, for 500 replicates.

| Sampling scheme | Summary statistics | Probability to find the true model |  |
| --- | --- | --- | --- |
|  |  | Without host alternation | With host alternation |
| Amance | Complete | 0.64 | 0.78 |
|  | Wrap-up | 0.67 | 0.81 |
| Grand Est | Complete | 0.73 | 0.81 |
|  | Wrap-up | 0.79 | 0.88 |

**Table 6:** Accuracy of the parameter estimation on simulated populations sampled on S, R and A compartments for the life cycle with host alternation, depending on the summary statistics considered. The sampling schemes of the simulation data sets match those of the two the empirical data sets: Amance location and Grand Est region. Numbers represent r-squared of the linear regressions between simulated and estimated parameters. The parameter identifiability procedure is based on a leave-one-out cross-validation with the neural networks regression method and tolerance parameter set at 0.01, for 200 replicates.

| Sampling scheme | Summary statistics | *propR* | *mig* | *f<sub>avr</sub>* | *r* | *K* | $\tau$ |
| --- | --- | --- | --- | --- | --- | --- | --- |
| Amance | Complete | 0.90 | 0.27 | 0.71 | 0.68 | 0.83 | 0.58 |
|  | Wrap-up | 0.90 | 0.39 | 0.77 | 0.79 | 0.89 | 0.67 |
| Grand Est | Complete | 0.91 | 0.31 | 0.71 | 0.68 | 0.81 | 0.68 |
|  | Wrap-up | 0.90 | 0.37 | 0.77 | 0.72 | 0.85 | 0.67 |

#### 3.3.2 Model choice

For the two empirical data sets, we infer the life cycle with host alternation with probability 100% from Complete summary statistics and the life cycle without host alternation with probabilities 81% from Wrap-up summary statistics (Table 7). However, in all cases, the results of the goodness-of-fit tests do not allow to significantly reject either of the life cycles, irrespective of the data set considered and the summary statistics used. Therefore, we do not have enough information to properly infer the pathogen life cycle. This is consistent with the coordinates of the data sets on the PCA analyses (Figure 4), which are located in the overlap area of the two models. This explains the weak identifiability of the life cycle and the impossibility of significantly rejecting either life cycle. Therefore, we have to rely on biological insights to determine the life cycle.

**Table 7:** Posterior model probabilities and goodness of fit for two data sets: in Amance location and the Grand Est region, depending on the summary statistics. With and Without stand for life cycles with and without host alternation, respectively. *P −values* < 0.01 are considered significant.

| Sampling scheme | Summary statistics | Posterior probability | | Goodness of fit ($P$-value) | |
| --- | --- | --- | --- | --- | --- |
|  |  | Without | With | Without | With |
| Amance | Complete | 0.00 | 1.00 | 0.069 | 0.045 |
|  | Wrap-up | 1.00 | 0.00 | 0.048 | 0.030 |
| Grand Est | Complete | 0.00 | 1.00 | 0.099 | 0.062 |
|  | Wrap-up | 0.81 | 0.19 | 0.092 | 0.064 |

**Figure 4:**
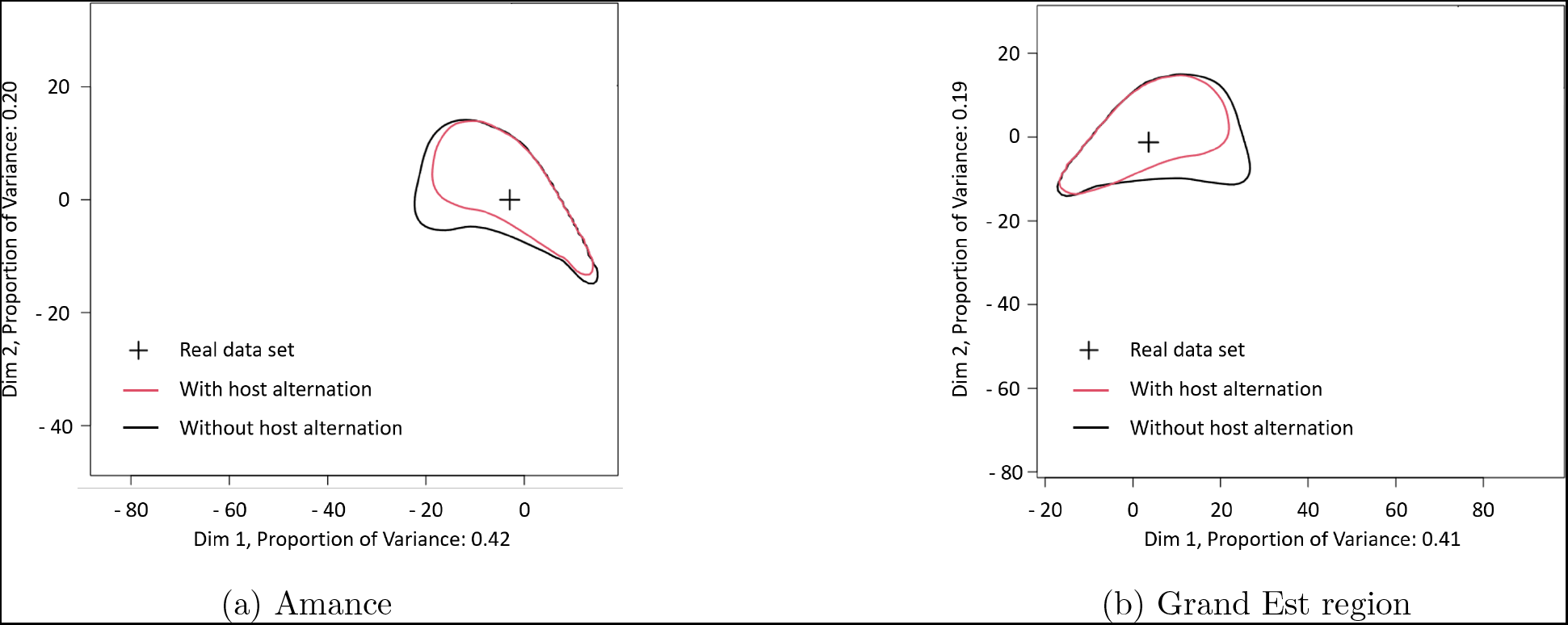
Principal component analyses (95% envelope) of simulations for each model: with and without host alternation, depending on the data set considered. Simulations were performed with Complete summary statistics. The black crosses correspond to the coordinates of the actual data sets.

#### 3.3.3 Parameter estimation

Knowing that the poplar rust pathogen alternates on larch to perform its sexual reproduction, we compute the parameter estimations under the model with host alternation.

Contrary to the theoretical analysis of parameter identifiability, Complete summary statistics perform better than Wrap-up summary statistics for the analysis of both empirical data sets. For most parameters, especially those that are well identified, the posterior distribution is more narrow and the difference between the posterior density versus the prior density is more pronounced (Figures 5, S1). Therefore, in the following, we focus on the parameter estimation from Complete summary statistics. As expected from the theoretical identifiability, narrow posterior distributions allow a confident inference of parameters *propR* and *K*. The strength of inference of parameters *f_avr_* and *τ* is more limited, and the posteriors for parameters *mig* and *r* are not much informative compared to their prior distributions. From the mode of each posterior distribution (Table 8), we infer a high proportion of resistant poplars at the time of resistance overcoming: *propR* = 0.78 and *propR* = 0.84 for Amance and the Grand Est region data sets respectively. The inferred population sizes are in the order of a thousand (mode values *K* = 1, 523 and 1, 356 for Amance and the Grand Est region, respectively). We infer an initial proportion of virulent alleles *f_avr_* in the pathogen population of 9% in Amance, and 6% in the Grand Est region, but with a large credible interval (range of values for which the posterior distribution is higher than the prior distribution), between 5% and 15%. The annual mortality rate *τ* during the annual migration preceding sexual reproduction is inferred to close values of 0.60 and 0.58 in Amance and in the Grand region, respectively. The migration rate *mig* and growth rate *r* are badly estimated (*mig* would range from 0.02 to 0.10, and *r* would be above 1.5).

**Table 8:** Statistical summary of the inference of the parameters for the life cycle with host alternation, depending on the data set.

| Data set | Parameter | $q - 2.5\%$ | median | mean | mode | $q - 97.5\%$ |
| --- | --- | --- | --- | --- | --- | --- |
| Amance | $propR$ | 0.49 | 0.81 | 0.78 | **0.88** | 0.94 |
| | $mig$ | 0.01 | 0.06 | 0.06 | **0.05** | 0.10 |
| | $f_{avr}$ | 0.03 | 0.11 | 0.11 | **0.09** | 0.19 |
| | $r$ | 1.31 | 1.69 | 1.68 | **1.92** | 1.98 |
| | $K$ | 1,205 | 2,024 | 2,561 | **1,374** | 6,414 |
| | $\tau$ | 0.51 | 0.66 | 0.67 | **0.58** | 0.88 |
| Grand Est | $propR$ | 0.42 | 0.77 | 0.74 | **0.80** | 0.94 |
| | $mig$ | 0.01 | 0.04 | 0.05 | **0.04** | 0.10 |
| | $f_{avr}$ | 0.02 | 0.08 | 0.09 | **0.06** | 0.19 |
| | $r$ | 1.31 | 1.69 | 1.67 | **1.80** | 1.96 |
| | $K$ | 768 | 2,206 | 3,108 | **1,273** | 10,146 |
| | $\tau$ | 0.51 | 0.64 | 0.66 | **0.58** | 0.88 |

**Figure 5:**
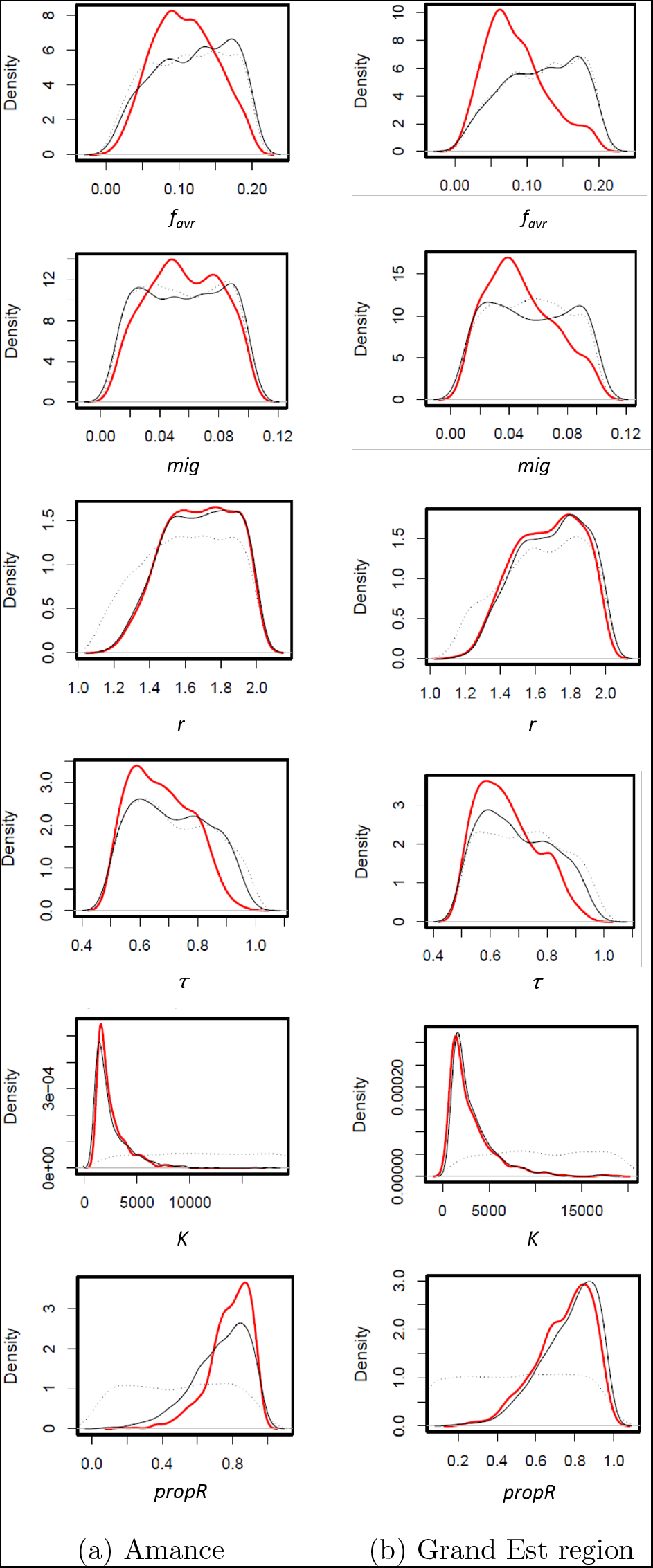
Posterior distributions of parameters, with Complete summary statistics, for the data sets in Amance location (on S and A) and the Grand Est region (on S, R and A). Dashed lines correspond to the prior distributions, black lines correspond to posterior distributions given by the rejection method, red lines correspond to the posterior distributions given by the neuralnet method.

## 4 Discussion

### Time samples unravel rapid adaptation

In this paper, we develop an original methodology to infer demographic scenarios and parameter values in the case of rapid adaptation. We base our inference framework on the use of time series data to grasp changes in population structure over time. We employ a forward population genetics model to simulate the summary statistics used for ABC inference. As such, our methodology contrasts with the most recent developments in population genetics inference that are based on the coalescent theory. Theoretical investigations demonstrate high accuracy in both model and parameter estimations. Last, our inferential framework is successfully applied to a case study of rapid adaptation in a plant pathogen following a resistance overcoming event with sound estimates of key demographic parameters.

### Too many summary statistics lead to model overfitting

The main originality of our method is to take into account the time series, compared to the high amount of studies that focus on a single time point. To do so, we adapt the classical ABC protocols of using summary statistics. We compare two methods, either keeping the complete sequence of statistics over all time samples or wrapping up the information into the mean, variance, minimum and maximum values for each summary statistic. We show in the theoretical cross-validation procedures that Wrap-up summary statistics outperform Complete summary statistics. This result may appear counter-intuitive. However too much and redundant information can lead to a decrease in statistical power. Since many statistics can be used in ABC, there is a certain curse of dimensionality, early on identified. This generates overfitting that may affect the accuracy of model and parameter estimation. This is especially acute when several summary statistics correlate with the same model parameters, inflating the heterogeneous accuracy inference of some parameters compared to others. A suggested solution is to perform first a dimension reduction of the statistics space before performing the ABC estimations. In our case, we have few genetic indices and few parameters, so we do not perform such reduction of dimensionality beforehand. We want to keep the full time series to avoid loss of temporal information, but in doing so we inflate the number of summary statistics which is likely to result in temporal auto-correlation causing overfitting. In support of this hypothesis, the discrepancy in parameter identifiability between Complete and Wrap-up statistics tends to decrease when we apply a thinning in the time series consisting in keeping only one fifth of the temporal samples. A way to keep the maximum information while avoiding the redundancy that causes overfitting would be to summarise the time series in a different manner. For example, we may fit a function describing the temporal changes and use regression coefficients as summary statistics. However, this method requires that all dynamics be correctly fitted by a similar function (with the same number of regression coefficients). In this attempt, we try to fit quadratic polynomes, but the shape of the temporal variation varied too much among population genetic indices to provide conclusive summary statistics (data not shown). Moreover, such fit cannot take into account stochastic variation due to drift. Another way to improve the inferential framework would be to avoid the use of population genetics indices as summary statistics and base the inference method directly on the evolution of genotypic frequencies, or a less condensed type of information such as site frequency spectra (SFS) across time samples. However this method may be more computationally intensive (in particular for storing the results over a long time series), and the SFS cannot be computed from microsatellite markers. Moreover, the resulting increase in the number of summary statistics could also lead to high dimensionality of the data.

### Case study estimates match the biology of the poplar rust system

By applying this method on empirical data sets, we infer accurately three parameters: *propR*, *K* and *f_avr_*. The estimated values make sense with regard to our knowledge of this event of resistance overcoming. We infer a very large proportion of resistant poplars (80%), which is consistent with our knowledge of poplar plantations before the RMlp7 resistance overcoming. Indeed, the resistant poplar cultivar ‘Beaupré’ that bears resistance RMlp7 was widely planted at the time of resistance overcoming and represented up to 80% of poplar cultivar sales in 1996 (data from the French Ministry of Agriculture, QUAEChap27Fabre). This very high proportion of resistant poplars in the landscape exerted a strong selection pressure and accounts for the changes in the genetic structure of a pathogen population over time (Persoons et al., 2017). We infer a population size of around one thousand individuals. This population size does not represent the number of individuals actually multiplying in the population, but those that effectively contribute to the observed genetic variability. This order of magnitude is consistent with a previous estimate based on a coalescent analysis (Persoons, 2015). We estimate an initial proportion of virulent alleles in the population between 5% and 10%. This range is consistent with the genetic characterisation at the selected locus in *M. larici populina* (Louet et al., 2021). Louet et al. (2021) highlighted that the alleles conferring virulence pre-existed in the pathogen population long before the resistance overcoming, at a frequency of 21%, on average, between 1989 and 1993. The proportion of virulent alleles in the population can strongly fluctuate during the years preceding resistance overcoming (Saubin et al., 2021). Such fluctuations are indeed observed in the data from Louet et al. (2021), with a standard deviation in the proportion of virulent alleles of 0.12 between 1989 and 1993. Our estimation is therefore consistent with the empirical data considering such large fluctuations from year to year. This level of standing genetic variation is not negligible in view of the adaptive potential it brings to the pathogen population.

The fact that some parameters are not well estimated is not caused by a limited amount of data, but rather by a strong assumption in our modelling framework. Indeed, we assume three compartments only in the landscape and neglect the more complex spatial structures that can be encountered in agricultural landscapes. In such a non-spatial system, it may be especially difficult to disentangle the genetic effects of growth, mortality and migration rates. We believe that increasing the spatial complexity of the model would help disentangle these three parameters. However, this would also highly increase the dimensions of summary statistics, which can lead to a decrease in the statistical accuracy of the ABC.

Despite the precautions taken when evaluating the theoretical accuracy of our method, we observe that the accuracy of our ABC estimation differs between the theoretical and empirical results. We obtain better theoretical results with Wrap-up summary statistics than with Complete summary statistics. However, Complete summary statistics lead to more accurate parameter inferences from empirical data sets. We believe that this is due to increased variability in population genetics indices in the empirical data compared to the simulated data. Many sources of biological stochasticity are not accounted for in our modelling assumptions. This stochasticity results in stronger local extremes, which are captured by the minimum and maximum values in Wrap-up summary statistics. Thus, taking into account all the available information of the empirical time series through Complete summary statistics can reduce the impact of this additional stochasticity, and improve the parameter estimations. This discrepancy between our application to empirical data and theoretical findings from simulated data can also originate from strong modelling assumptions (like the non-spatial model) that do not perfectly reflect the biological system.

### Methodological guideline

We recommend using a sampling scheme with regular time points to capture sufficient information in the genetic evolution of populations. Contrary to the current practice, we show that for this framework it is more efficient to sample more populations in time but on a single host than to focus on few temporal samples but on several hosts. The sampling scheme with the first and the last time points, even if less informative than a more regular sampling, still allows to correctly infer some of the model parameters. In particular, the initial frequency of virulent alleles in the population is very well inferred with this sampling scheme because the first population in time is the most important to identify the initial genetic composition of the population.

### Future directions: application for genome scans of rapid adaptation

Neutral genetic data from microsatellite markers contain sufficient information to infer parameters of resistance overcoming from time samples. However, we could still gain in accuracy by taking genomic data into account. The integration of genomic data would allow the calculation of a wider range of summary statistics, which could increase the accuracy of inferences. Fitting demographic models is a prerequisite for detecting selection, but it can be difficult to do in practice and is often inaccurate (Hoban et al., 2016). The addition of genomic data to the described framework would enable to focus on areas under selection, detect sweeps (Messer and Petrov, 2013; Foll et al., 2015) and calculate their age. Determining the evolution of neutral loci through such a selection event may allow, by comparison, to identify selected loci implied in rapid adaptation.

## Acknowledgements

We warmly thank Emma Chavan for her help with the microsatellite analyses. This work was supported by grants from the French National Research Agency (ANR-18-CE32-0001, Clonix2D project; ANR-11-LABX-0002-01, Cluster of Excellence Arbre) and from the Metaprogram SuMCrop of the National Research Institute for Agriculture, Food and the Environment (INRAE, Opiniâtres project). Méline Saubin was supported by a PhD fellowship from INRAE and the French National Research Agency (ANR-18-CE32-0001, Clonix2D project). Méline Saubin obtained an international mobility grant BayFrance as part of a Franco-Bavarian cooperation project, to work for one month in Aurélien Tellier’s lab (Grant Number FK21 2020).

## Data accessibility

Python codes for model simulations, R scripts for statistical analyses and data for the biological application will be made available on Zenodo repository at the time of publication.

## Author contributions

Méline Saubin, Aurélien Tellier and Fabien Halkett conceived and designed the study. Axelle Andrieux performed the additionnal genotyping. Méline Saubin and Solenn Stoeckel produced the code and ran the simulations. Méline Saubin, Aurélien Tellier and Fabien Halkett analysed the data and prepared the manuscript. All authors revised and approved the manuscript.

## Appendix

## A Effect of sampling rarefaction applied to compartments and time series

To investigate the effect of a reduced amount of information, we perform ABC analyses based on rarefied sampling scheme. Cross-validation procedures for the model choice (Table S1) and parameter estimation (Table S2) are performed as presented in Section 2.3.

Two types of rarefaction are applied, affecting the time series and the range of compartments considered.

Concerning the time series, in addition to the full time series, we consider (1) a thinning that keeps samplings every five years for a total of eight time samples and (2) only the first and last time samples. For the latter temporal rarefaction, it is not possible to compare the two types of summary statistic since only two generations are considered.

Concerning rarefaction of the sampled compartment (S, R or A) we apply different rarefaction types depending on the cross-validation procedure. The model choice procedure is based on: (1) Populations sampled on both S and R; (2) Populations sampled on S only. Populations sampled on A are not used for the model choice because this compartment only exists for the life cycle with host alternation. The parameter estimation procedure focus on the life cycle with host alternation and is based on: (1) Populations on S, R, and A; (2) Populations on S and R; (3) Populations on S and A; (4) Populations on S only.

**Table S1:**
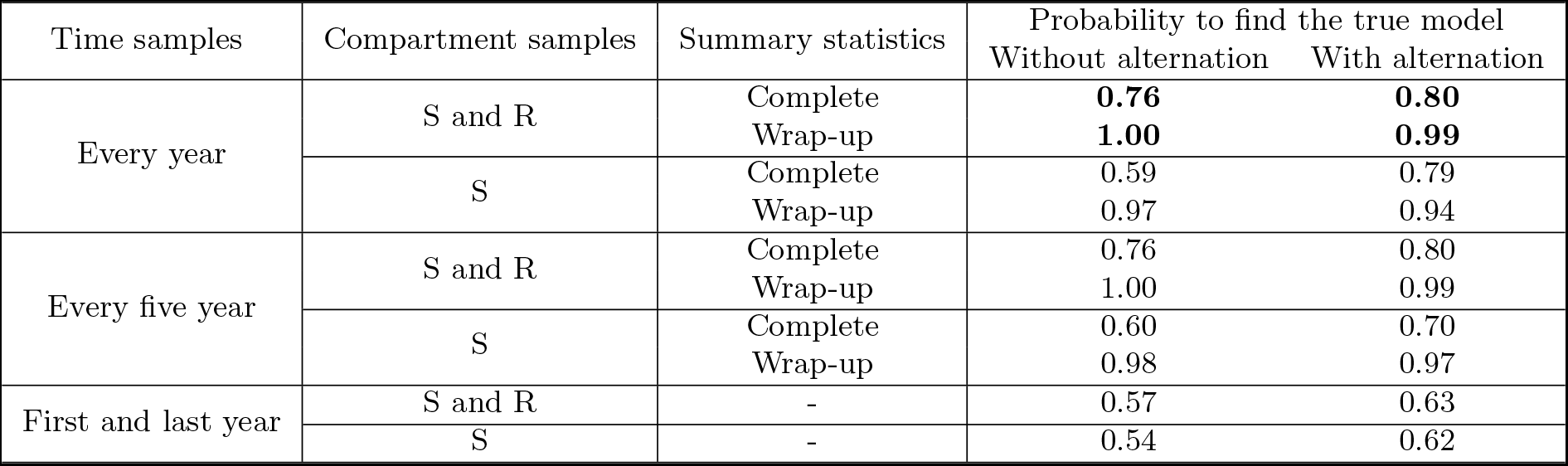
Accuracy of model selection depending on the summary statistics and the sampling rarefaction considered. The model choice procedure is based on leave-one-out cross-validations with a weighted multinomial logistic regression computed with tolerance parameter set at 0.01, for 500 replicates. Bold values represent the values reported in the main text.

**Table S2:** Accuracy of the parameter estimation depending on the summary statistics considered and the type of rarefaction, for simulated populations with host alternation. Data represent r-squared of the linear regressions between simulated and estimated parameters. The parameters identifiability procedure is based on a leave-one-out cross-validation with the neural networks regression method and tolerance parameter set at 0.01, for 200 replicates. Bold values represent the values already presented in the main text.

| Time samples | Compartment samples | Summary statistics | $propR$ | $mig$ | $f_{avr}$ | $r$ | $K$ | $\tau$ |
| --- | --- | --- | --- | --- | --- | --- | --- | --- |
| Every year | S, R and A | Complete | **0.91** | **0.41** | **0.68** | **0.73** | **0.81** | **0.58** |
|  |  | Wrap-up | **0.94** | **0.60** | **0.86** | **0.85** | **0.92** | **0.79** |
|  | S and R | Complete | 0.80 | 0.33 | 0.75 | 0.78 | 0.79 | 0.56 |
|  |  | Wrap-up | 0.94 | 0.70 | 0.85 | 0.83 | 0.90 | 0.77 |
|  | S and A | Complete | 0.90 | 0.23 | 0.67 | 0.65 | 0.79 | 0.62 |
|  |  | Wrap-up | 0.94 | 0.43 | 0.83 | 0.79 | 0.90 | 0.78 |
|  | S | Complete | 0.89 | 0.31 | 0.70 | 0.63 | 0.85 | 0.64 |
|  |  | Wrap-up | 0.91 | 0.51 | 0.81 | 0.73 | 0.88 | 0.73 |
| Every five year | S, R and A | Complete | 0.89 | 0.31 | 0.73 | 0.68 | 0.80 | 0.68 |
|  |  | Wrap-up | 0.90 | 0.46 | 0.82 | 0.68 | 0.85 | 0.69 |
|  | S and R | Complete | 0.88 | 0.33 | 0.78 | 0.60 | 0.82 | 0.65 |
|  |  | Wrap-up | 0.92 | 0.31 | 0.85 | 0.68 | 0.86 | 0.69 |
|  | S and A | Complete | 0.91 | 0.32 | 0.78 | 0.67 | 0.80 | 0.72 |
|  |  | Wrap-up | 0.90 | 0.27 | 0.79 | 0.65 | 0.84 | 0.69 |
|  | S | Complete | 0.87 | 0.23 | 0.76 | 0.62 | 0.82 | 0.63 |
|  |  | Wrap-up | 0.89 | 0.46 | 0.81 | 0.61 | 0.85 | 0.65 |
| First and last year | S, R and A | - | 0.87 | 0.45 | 0.85 | 0.52 | 0.83 | 0.64 |
|  | S and R | - | 0.81 | 0.46 | 0.83 | 0.30 | 0.81 | 0.58 |
|  | S and A | - | 0.78 | 0.26 | 0.83 | 0.54 | 0.82 | 0.68 |
|  | S | - | 0.82 | 0.24 | 0.74 | 0.34 | 0.79 | 0.51 |

## B Case study: parameters’ inference with Wrap-up summary statistics

**Figure S1:**
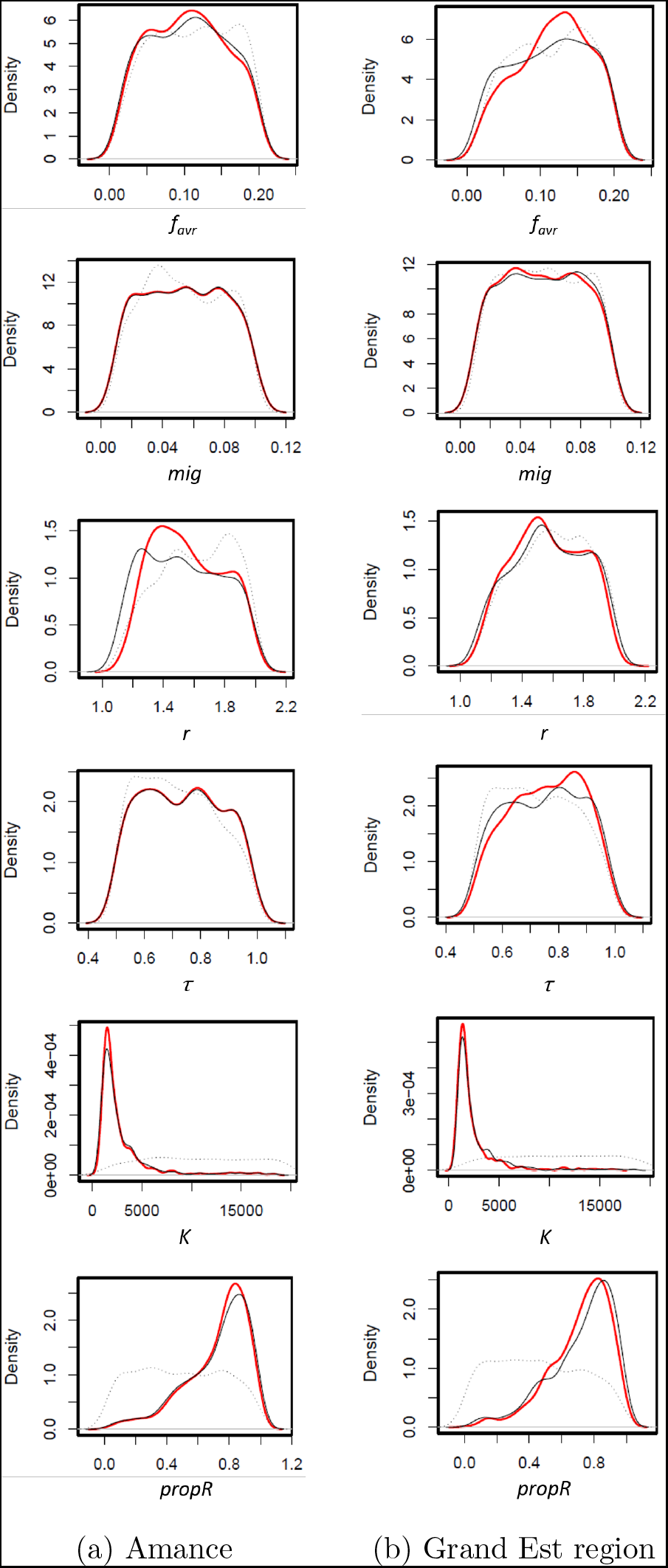
Posterior distributions of parameters, with Wrap-up summary statistics, for the datasets in Amance location (on S and A) and the Grand Est region (on S, R and A). Dashed lines correspond to the prior distribution, black lines correspond to posterior distributions given by the rejection method, red lines correspond to the posterior distributions given by the neuralnet method.

